# TET1 controls *Cxcl1* induction by DNA demethylation and promotes neutrophil recruitment during acute lung injury

**DOI:** 10.1101/2021.09.07.459280

**Authors:** Kathleen M. Yee, Richard W. Shuai, Bin Liu, Christian A. Huynh, Chao Niu, Hailey R. Lee, Min S. Lee, Jirui Wen, Jian Zou, Jiang Wu, Ke Shuai

## Abstract

Neutrophils are rapidly recruited from the peripheral blood to the inflammatory site to initiate inflammatory response against pathogenic infections. The process to recruit neutrophils must be properly regulated since the abnormal accumulation of neutrophils can cause organ damage and dysfunction. The acute respiratory distress syndrome (ARDS)/acute lung injury (ALI) is a common cause of respiratory failure that is characterized by the infiltration of neutrophils and epithelial integrity disruption. Indeed, recent studies suggest a pathogenic role of neutrophils in the clinic severity of the coronavirus disease 2019 (COVID-19) ARDS. The chemokine CXCL1, which is rapidly induced by inflammatory stimuli, plays a key role in neutrophil influx during lung inflammation. The molecular basis of *Cxcl1* induction is not fully understood. Here we report that TET1, a member of the ten eleven translocation (TET) methylcytosine dioxygenase protein family, displays a striking specificity in the regulation of gene expression in macrophages. RNA sequencing (RNA-seq) analysis showed that *Tet1* disruption significantly altered the expression of only 48 genes that include *Cxcl1* and several other genes known to be important for cell migration and trafficking in bone marrow derived macrophages (BMDMs) in response to LPS stimulation. TET1 regulates the induction of *Cxcl1* by facilitating the DNA demethylation of the *Cxcl1* promoter. In *Tet1*^−/−^ mice, the induction of *Cxcl1* was suppressed, resulting in defective neutrophil recruitment to the lung during LPS-induced acute lung injury. Our results identify a novel epigenetic mechanism that selectively controls *Cxcl1* induction and neutrophil recruitment during acute lung injury.

**Key Points:**

- TET1 has a striking specificity in macrophage gene regulation and controls *Cxcl1* induction by inflammatory stimuli via DNA demethylation
- Neutrophil recruitment is defective in *Tet1* deficient mice during acute lung injury

## Introduction

The acute respiratory distress syndrome (ARDS)/acute lung injury (ALI) is a common cause of respiratory failure that is associated with a high mortality, which ranges from 30% to 40% ^1^. The coronavirus disease 2019 (COVID-19) is caused by severe acute respiratory syndrome coronavirus 2 (SARS-CoV-2), and 10-15% of COVID-19 patients progress to ARDS ^2,3^. Neutrophils represent the first line of defense in response to invading pathogens and they are the key effector cells during tissue injury-induced inflammation ^4–6^. Upon pathogenic infections, neutrophils are rapidly recruited from the peripheral blood to the inflammatory site where they participate in the eradication of microbial infections. Individuals with a deficiency in neutrophils are more susceptible to infections. However, the process to recruit neutrophils to initiate inflammatory response must be properly regulated as the abnormal accumulation of neutrophils at the inflammatory site can cause organ damage and dysfunction ^5,7,8^. The role of neutrophils in ARDS/ALI is highlighted by their increased recruitment in the pulmonary capillary bed and into the alveolar space ^9^. Recent studies suggest a pathological role for neutrophils and neutrophil-derived neutrophil extracellular traps (NETs) in the clinic severity of COVID-19 ^10–14^. Understanding the mechanisms underlying neutrophil recruitment is important for the development of treatment strategies for ARDS/ALI.

The chemokine CXCL1 signals through its receptor CXCR2 to regulate cell migration and recruitment in immune defense, wound healing and tumorigenesis ^15–17^. Previous studies have demonstrated a key role of CXCL1 in neutrophil recruitment during lung inflammation ^18–22^. It has been well documented that there exists host difference in response to bacterial or viral infections, but the molecular basis involved remains to be poorly understood. Genetic or epigenetic factors may contribute to the differences in host response toward pathogens. In response to inflammatory stimuli, CXCL1 is strongly induced, and transcription factors NF-κB and STAT1 are shown to be involved in *Cxcl1* induction ^23^. The hypermethylation of *Cxcl1* promoter was observed in certain cancer cells ^24^. Whether epigenetic mechanism is involved in the transcriptional regulation of *Cxcl1* during inflammation is not known.

DNA modifications such as 5-hydroxymethylcytosine (5hmC) are catalyzed by the ten eleven translocation (TET) methylcytosine dioxygenase protein family, and TET proteins are known to participate in the DNA demethylation process ^25–32^. Gene targeting studies have uncovered a key role of TET proteins in multiple cellular processes ^33–38^. The role of TET proteins in immune regulation is best exemplified by the finding that TET2 acts to restrict the production of inflammatory cytokines such as IL-6 and IL-1β in macrophages ^38^, and the loss of TET2 results in clonal hematopoiesis and accelerates atherosclerosis ^39,40^. Surprisingly, despite the well established enzymatic role of TET2 in promoting DNA demethylation, it was found that TET2 functions as a transcriptional repressor to suppress inflammatory gene activation by recruiting histone deacetylases (HDACs), a mechanism that is independent of its activity in DNA demethylation ^38,39^. These findings have raised an interesting question on whether the TET-mediated DNA demethylation plays an important role in immune regulation.

In this report, we describe a highly specific role of TET1 in the regulation of *Cxcl1* induction in response to inflammatory stimuli such as LPS and TNFα. TET1 acts by regulating the DNA methylation of the *Cxcl1* promoter. We demonstrated that the TET1-CXCL1 regulatory pathway is important for the recruitment of neutrophils during acute lung injury.

## Materials and methods

### Animal treatment

The animal work conducted in this manuscript was approved by the Animal Research Committee of the University of California Los Angeles. *Tet1*^+/−^ mice in C57BL/6 129Sv genetic background ^41^ were purchased from the Jackson lab and have been backcrossed to C57 or 129 background for 10 generations. For acute lung injury experiment, WT and *Tet1*^−/−^ mice were intranasally injected with 0.5mg/kg LPS (E. coli 0111:B4, Sigma) for 4 hr.

### Primary cell cultures

Mouse bone marrow-derived macrophages (BMDMs) were prepared as previous described ^42^. Briefly, bone marrow was flushed from the lower leg bones of mice and subsequently treated with ACK buffer (0.15 M NH4Cl, 10 mM KHCO3, 0.1 mM EDTA) to lyse red blood cells. Then 3×10^5^ cells were seeded per 35 mm well with media containing DMEM, 10% FBS, 1% penicillin-streptomycin and 30% L929 conditioned media. Media was changed at either day 3 or day 4. At day 7 cells were used for experiments.

For preparing mouse embryonic fibroblast cells (MEF), mouse embryos were collected around embryonic day 14. Briefly, visible organs such as the brain, liver, and spleen were removed, and the leftover tissue was quickly minced and incubated in 0.25% trypsin-EDTA for 2 hr at 37°C. Cells were then washed with media and seeded in DMEM with 10% FBS, 1% penicillin/streptomycin and 0.055 mM β-mercaptoethanol. Every three days cells were passaged. Starting at passage 2, cells were used for experiments.

### RNA sequencing and analysis

Bone marrow derived macrophages from WT and *Tet1*^−/−^ mice were untreated or treated with LPS for 4 hr. Total RNA was collected and prepared with STAT60 (Tel-Test, Inc.) according to manufacturer’s protocol, and was then column purified using RNA Clean and Concentrator (Zymo Research), according to manufacturer’s protocol. Library preparation, amplification, and RNA sequencing (HiSeq3000) were performed by the UCLA Technology Center for Genomics and Bioinformatics (TCGB) core.

For RNAseq analysis, the raw sequencing reads (FASTQ files) were first preprocessed to remove low-quality reads and bases. The data was screened for contamination with FASTQScreen, and the quality of the FASTQ sequencing data was assessed with FastQC ^43^ (http://www.bioinformatics.babraham.ac.uk/projects/fastqc/). Alignment and differential gene expression analysis were performed using a Snakemake workflow ^44^. The reads were aligned to the mm10 reference genome from the Ensembl genome database and counted using STAR ^45^. Normalization of raw counts and detection of differentially expressed genes (DEGs) were performed with the DEG elimination strategy (DEGES) from the TCC package using the TMM-(edgeR-TMM)_3_ pipeline followed by DEG identification with edgeR ^46,47^. Adjusted p-values were determined using the Benjamini–Hochberg procedure in identification. Heatmaps were generated using the pheatmap R package (https://cran.r-project.org/package=pheatmap). For display on heatmaps, normalized counts were converted to a log_2_ scale, and the z-scores were calculated using the means and standard deviations across the samples for each gene.

### Promoter Methylation Analysis

Genomic DNA (gDNA) was converted with sodium bisulfite using the EZ Methylation Gold Kit (Zymo Research) according to manufacturers protocol. *Cxcl1* promoter regions #1 and #2 were PCR amplified from sodium bisulfite converted gDNA using ZymoTaq Polymerase (Zymo Research) and purified by gel extraction using the QIAquick Gel Extraction Kit (Qiagen). These purified PCR products were then cloned into pCR4-TOPO using the TOPO-TA Cloning Kit (Thermo Fisher Scientific) and transformed into chemically competent cells. Minipreps were screened for correctly sized inserts and positive clones were sent for sequencing.

The following primers were used to amplify the *Cxcl1* promoter region from converted gDNA:

*Cxcl1* region 1 methylation-5’: GGGGTAAAAAGTAAAAATTTTAGAGTTTAAT
*Cxcl1* region 1 methylation-3’: TCCCAACCATAATCTACAAACACTAACAAC
*Cxcl1* region 2 methylation-5’: AGTAGGGTAGGGATGTTTTAGGAATTAT
*Cxcl1* region 2 methylation-3’: TTAAACTCTAAAATTTTTACTTTTTACCCC

### Luciferase Reporter Analysis

*Cxcl1* promoter region from nucleotide −157 to +283 was cloned into pGL3-basic (Promega) through KpnI and BglII cloning sites using the following primers:

*Cxcl1*_promoter 5’: CGGTACCCAAAAAGCAAAAATCCCAGAGT
*Cxcl1*_promoter 3’: CAGAGATCTCTCTGGATGTTCTTGAGGTGAA

To create artificially methylated *Cxcl1* promoter in pGL3 expression vector, the PCR-amplified *Cxcl1* promoter fragment was digested with KpnI and BglII (New England Biolabs) and then purified by gel extraction using the QIAquick Gel Extraction Kit (Qiagen). The purified *Cxcl1* promoter fragment was equally split into two groups: one treated with M.SssI (New England Biolabs) and the other without M.SssI (“mock”) according to manufacturers’ protocol. The two treatment groups were then column purified and ligated into pGL3 basic using T4 ligase (Thermo Fisher Scientific). Luciferase promoter activity was determined using the Dual-Luciferase Reporter Assay System (Promega) according to manufacturer’s protocol. 293T cells were transfected by calcium phosphate method with the various pGL3 expression vectors along with an equal amount of Renilla plasmid. The next day, cells were treated with or without TNF⍰ (20 ng/ml) for 5 h to induce *Cxcl1* expression. Firefly luciferase values were normalized by Renilla luciferase expression.

### Flow Cytometry Analysis

To collect bronchoalveolar lavage (BAL) fluids, mouse ribs were carefully removed to expose the lung cavity and neck tissues removed to expose the trachea of euthanized mice. A 22 G catheter was then inserted into the trachea. Using a 1 ml syringe, 0.75 ml of cold PBS was injected into the lungs via the catheter and the fluid aspirated out and collected. This was repeated four more times, for a total volume of roughly 3.5 ml. Single cell suspensions of the BAL fluids were stained with various combinations of surface markers followed by flow cytometry analysis using a FACSCalibur or FACScanX (Becton Dickinson, BD). Data were analyzed using the FlowJo software. Neutrophils, macrophages, and eosinophils were identified on the Flow as previously described ^48,49^. Specifically, neutrophils were gated as CD45^+^CD11b^+^Ly6G^+^ cells, macrophages were identified as CD45^+^CD11c^+^SiglecF^+^ population of cells, and eosinophils were defined as CD45^+^CD11c^low/−^SiglecF^+^ cells.

### Quantitative real time polymerase chain reaction (Q-PCR) analysis

Quantitative real time polymerase chain reaction (Q-PCR) analyses were performed as described ^42^. Briefly, cDNA prepared using the iScript cDNA Synthesis kit (Bio-Rad) were analyzed by Q-PCR using the CFX-96 Real-Time Detection System (Bio-Rad). See Supplementary Information for primer sequences of genes analyzed by Q-PCR.

### ELISA assays

Enzyme-linked immunosorbent assay (ELISA) kits for CXCL1 (Peprotech) and TNFα (R&D) were used to measure CXCL1 and TNFα protein levels in BAL fluid following the manufacturer’s instructions.

## Results

### Systematic analysis of the role of TET1 in LPS-triggered gene induction by RNA-seq

To examine if TET1 is regulated by inflammatory stimuli, WT and *Tet1*^−/−^ BMDMs were treated with a variety of Toll-like receptor (TLR) ligands followed by Q-PCR analysis. LPS stimulation strongly increased *Tet1* mRNA in WT but not *Tet1*^−/−^ BMDMs. *Tet1* was also induced in response to TLR2 ligand Zymosan and TLR3 agonist Poly(I:C), but not IFNγ or IL-4 (Fig. 1A). These results suggest that TET1 may play a role in TLR signaling.

**Figure 1.**
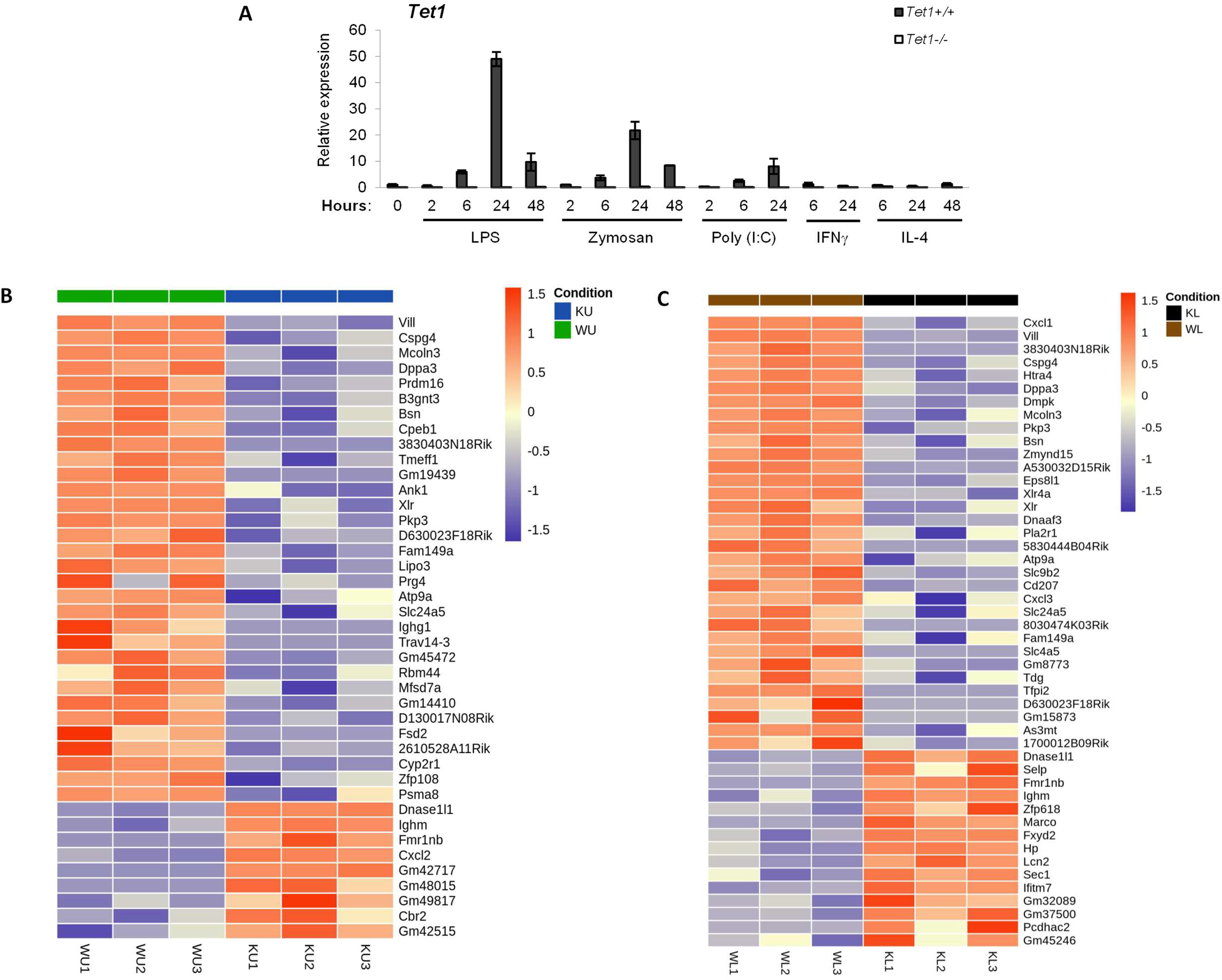
RNA-seq analysis of WT and *Tet1*^−/−^ BMDMs upon LPS stimulation. (A) TET1 is induced by TLR signaling. Quantitative real time polymerase chain reaction (Q-PCR) analysis of *Tet1* expression in WT and *Tet1*^−/−^ BMDMs in response to LPS (10ug/ml), Zymosan (50 ug/ml), Poly (I:C) (20 ug/ml), IFNγ (5 ng/ml), and IL-4 (10 ng/ml) for various time periods as indicated. (B) and (C) Heat maps of the differentially expressed genes (fold change > 2; adjusted P value < 0.05) in untreated WT (WU) vs untreated *Tet1*^−/−^ (KU) BMDMs (B) and WT (WL) vs *Tet1*^−/−^ (KL) BMDMs treated with LPS for 4 hr (C). WU: wild type untreated; KU: knockout untreated; WL: wild type treated with LPS; KL: knockout treated with LPS. WU1-3 and KU1-3 represent independent biological samples.

To examine the role of TET1 in the LPS-induced gene regulation, we performed a non-biased systematic study of gene profiling of WT and *Tet1*^−/−^ BMDMs untreated or treated with LPS for 4 hr by RNA-seq analysis. Surprisingly, as shown in Fig. 1B & 1C, the expression of only a very limited number of genes was significantly affected by *Tet1* deletion either constitutively or upon LPS stimulation (fold change >1.5, adjusted P value <0.05). *Tet1* deletion caused the significant suppression of 32 and 33 genes in untreated or LPS treated cells, respectively, while the removal of TET1 induced only 9 genes in untreated cells and 15 genes upon LPS stimulation.

The most significantly repressed gene caused by *Tet1* deletion upon LPS stimulation is *Cxcl1*, a chemokine known to regulate cell migration and recruitment ^15–17^ (Fig. 1C). In addition to *Cxcl1*, *Tet1* disruption also resulted in the altered expression of a number of genes known to regulate cell migration and trafficking under various physiological settings, including chondroitin sulfate proteoglycan 4 (*Cspg4*), cytoplasmic polyadenylation element-binding protein 1 (*Cpeb1*), serine protease HTRA4 (*Htra4*), proteoglycan 4 (*Prg4*), and N-acetyllactosaminide beta-1,3-N-acetylglucosaminyltransferase 3 (*B3gnt3*) ^50–55^. Thus, RNA-seq analysis revealed that TET1 displayed a striking specificity in gene regulation with the most recognizable preference for the regulation of genes involved in cell migration and trafficking.

### TET1 specifically regulates the induction of *Cxcl1* in multiple cell types

Next, we performed Q-PCR analysis to validate the findings of the RNA-seq studies. The induction of *Cxcl1* was defective in *Tet1*^−/−^ BMDMs in response to LPS stimulation for 2 hr, 6 hr or 24 hr, while the induction of other chemokine genes such as *Cxcl2* and *Cxcl10* was not affected (Fig. 2A). The removal of one copy of *Tet1* (*Tet1^+/−^*) showed only a modest reduction on *Cxcl1* expression as compared to the WT control, and the level of *Cxcl1* in *Tet1^+/−^* cells was at least 10xfold higher than that in *Tet1^−/−^* cells (supplementary Fig. S1).

**Figure 2.**
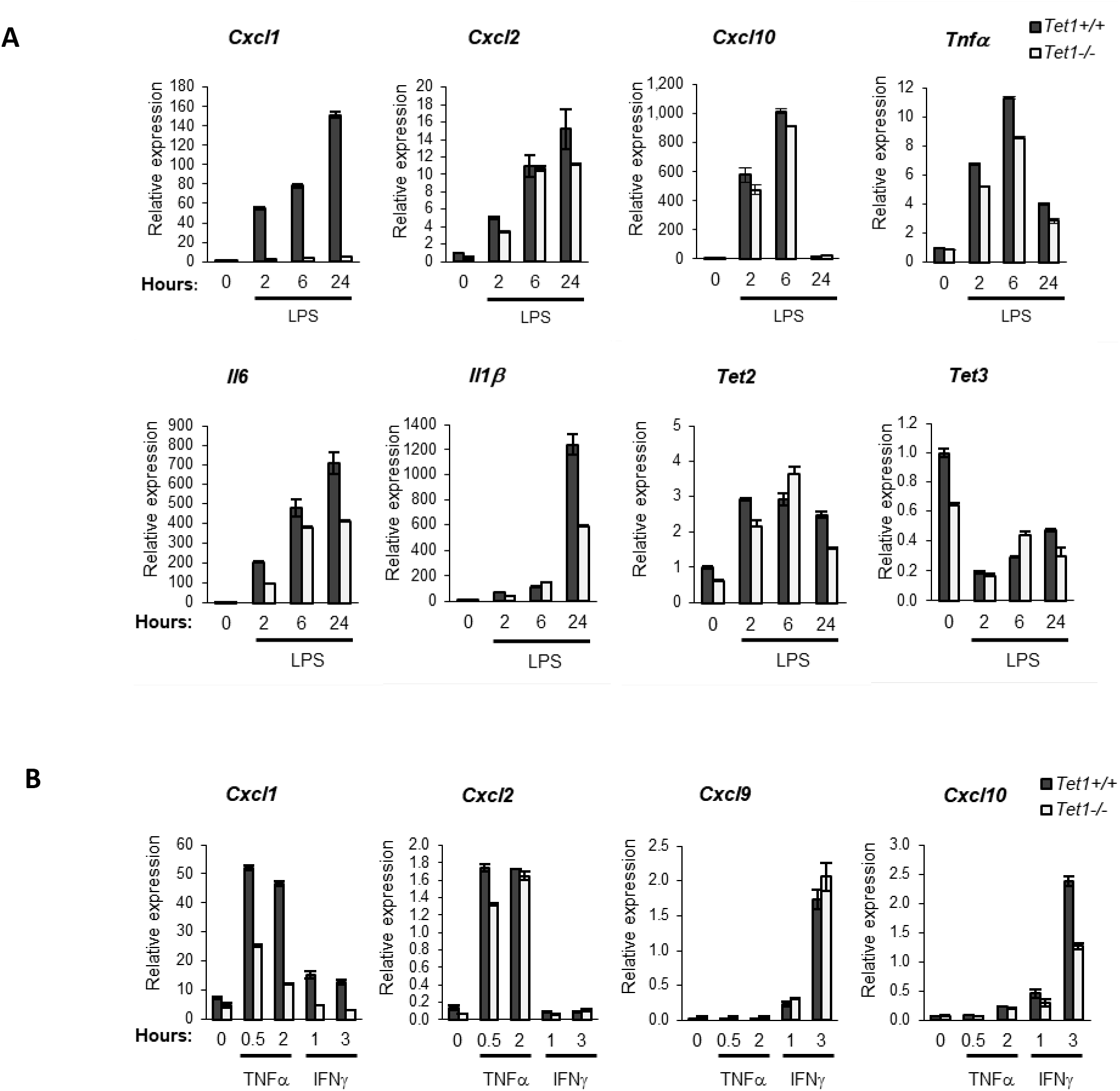
TET1 selectively regulates the expression of a subset of genes in response to inflammatory stimuli. (A) Q-PCR analysis of gene expression in WT and *Tet1*^−/−^ BMDMs treated with LPS (10 ng/ml) for 2 hr, 6 hr, and 24 hr as indicated. (B) Same as in (A) except RNA samples prepared from primary embryonic fibroblasts (MEFs) treated with TNFα (20 ng/ml) for 30 min or 2 hr or treated with IFNγ (5 ng/ml) for 1hr and 3 hr as indicated. Shown is the representative result from at least 3 independent experiments. Error bars represent SD.

Previous studies indicate that *Tet2* deletion in BMDMs altered the expression of over 450 genes, and caused the up-regulation of inflammatory genes such as *Il-6* and *Il1-β* ^40^. Consistent with the RNA-seq results, the induction of inflammatory cytokines such as *Il-6*, *Il-1β* or *Tnfα* was not significantly altered in *Tet1* deficient BMDMs in response to LPS stimulation for 2 hr or 6 hr. However, a prolonged treatment of *Tet1*^−/−^ BMDMs with LPS for 24 hr caused a modest repression of *Il-6* and *Il-1β*. The deletion of *Tet1* had no significant effect on the expression of *Tet2* or *Tet3* (Fig. 2A). Thus, TET1 and TET2 display opposing effects on inflammatory gene expression.

To further validate the specific effect of TET1 on *Cxcl1* expression, we performed Q-PCR analysis utilizing primary peritoneal macrophages isolated from wild type and *Tet1*^−/−^ mice or bone marrow derived dendritic cells (BMDCs) (Fig. S2). Again, the induction of *Cxcl1* was repressed in these cell types. These results support a key role of *Tet1* in the LPS-induced *Cxcl1* expression.

*Cxcl1* is known to be induced by other inflammatory cytokines such as TNFα. To test if the observed *Tet1*-*Cxcl1* regulation also operates in response to inflammatory stimuli other than LPS, we analyzed the effect of *Tet1* disruption on gene induction in response to TNFα or IFNγ in primary embryonic fibroblasts (MEFs) (Fig. 2B). *Cxcl1* was strongly induced by TNFα in WT but not *Tet1* deficient fibroblasts. Again, the induction of *Cxcl2* was not affected. *Cxcl1* was also induced by IFNγ in wild type MEFs, but not in *Tet1* deficient cells. In contrast, the induction of *Cxcl9* by IFNγ was not affected in the absence of TET1. The IFNγ-induced *Cxcl10* was modestly suppressed in *Tet1*^−/−^ MEFs (Fig. 2B). These results indicate that TET1 regulates the induction of *Cxcl1* in response to multiple inflammatory stimuli.

### TET1 regulates the promoter DNA methylation of *Cxcl1*

Results from Fig.1 and Fig. 2 indicate that the effect of TET1 in transcription is gene-specific in response to multiple stimuli. Although transcription factors such as STAT1 and NF-κB are known to be involved in the regulation of the expression of *Cxcl1* ^23^, it is unlikely that these transcription factors are directly involved in the observed specific regulation of *Cxcl1* induction since many well known targets of STAT1 and NF-κB were not affected in *Tet1* deficient cells. We hypothesized that TET1 might regulate *Cxcl1* expression through an epigenetic mechanism since TET1 is known to promote DNA demethylation.

TET proteins can promote DNA demethylation of CpG sites present in gene promoters. We identified two regions (Region #1 and Region #2, Fig. 3) within the *Cxcl1* promoter that contain potential CpG methylation sites. These two regions also contain potential NF-κB or STAT1 binding sequences (Fig. 3). We performed bisulfite-sequencing analysis and showed that these two regions were indeed enriched in CpG methylation in wild type BMDMs (Fig. 3 and Fig. S3). Interestingly, the extent of CpG methylation in Region #1 was clearly increased in *Tet1*-deficient BMDMs (Fig. 3). In Region #2, the extent of CpG site methylation was near saturation in wild type cells, and *Tet1* disruption did not show significant effect on DNA methylation in this region (Fig. S3). These results indicate that TET1 participate in the CpG site methylation in the region #1 of the *Cxcl1* gene promoter in BMDMs. Similar results were observed in the bisulfite-sequencing analysis of DNA samples from peritoneal macrophages of WT and *Tet1*^−/−^ mice (Fig.S4).

**Figure 3.**
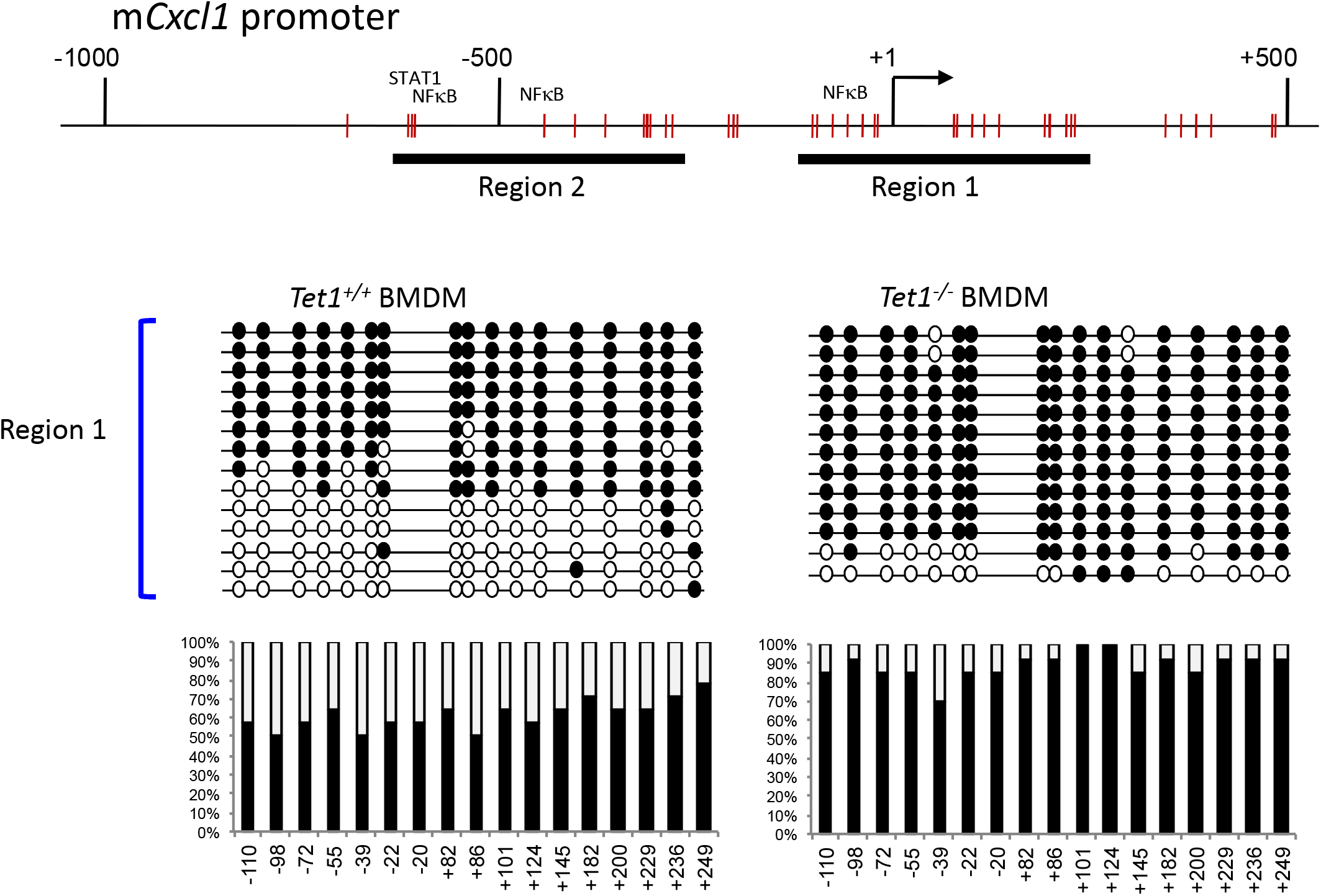
TET1 regulates the DNA methylation of the *Cxcl1* promoter in BMDMs. (A) Diagram of the murine *Cxcl1* promoter region approximately from −700 bp to +500 bp of the transcription start site. Short vertical lines represent potential CpG methylation sites. Regions #1 and #2 indicate two areas of interest for methylation analysis. (B) Methylation analysis of the *Cxcl1* promoter region#1 was performed by bisulfite conversion of genomic DNA from WT and *Tet1*^−/−^ BMDMs. The *x* axis represents the positions of the CpG sites relative to the transcription start site (+1); the *y* axis represents the percentage. Closed circle: methylated cytosine; Open circle: unmethylated cytosine.

To test if the TET1-mediated regulation of promoter CpG methylation also occurs in cells other than macrophages, we performed bisulfite-sequencing analysis of region #1 of the *Cxcl1* gene promoter using WT and *Tet1*^−/−^ primary embryonic fibroblasts (MEFs). In untreated wild type MEFs, the CpG site methylation in Region #1 was weak (10% to 30%) and was almost completely removed after TNFα stimulation (Fig. 4). In contrast, the CpG site methylation of Region #1 was found to be near 100% and remained to be very high (70% to 100%) after TNFα treatment in *Tet1*-deficient fibroblasts (Fig. 4). Taken together, our results suggest that TET1 controls the promoter DNA methylation of the *Cxcl1* gene in multiple cell types.

**Figure 4.**
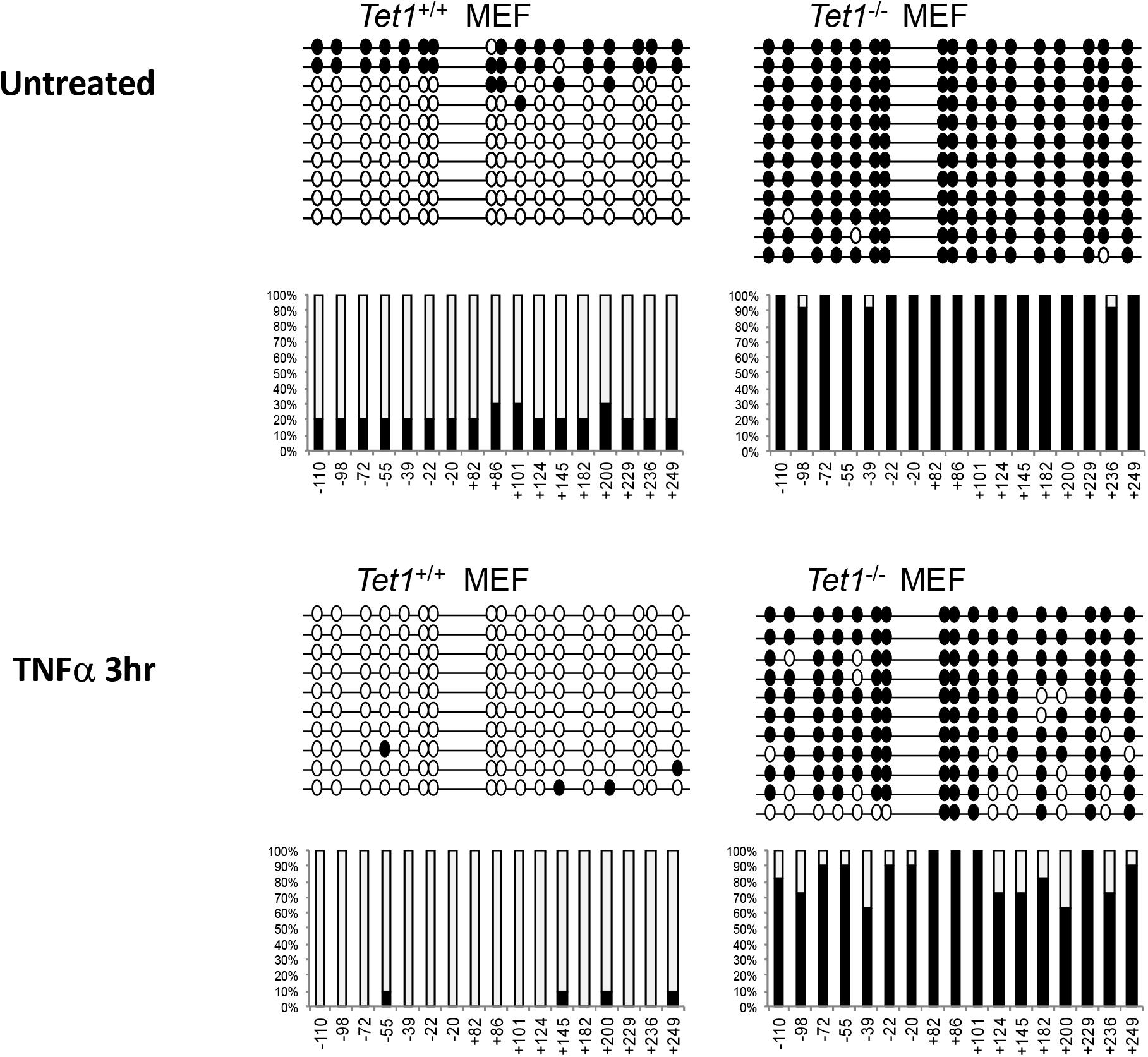
TET1 regulates the DNA methylation of the *Cxcl1* promoter in primary fibroblasts. Methylation analysis of the *Cxcl1* promoter region#1 was performed by bisulfite conversion of genomic DNA from WT and *Tet1*^−/−^ primary embryonic fibroblasts (MEFs) untreated or treated with TNFα (20 ng/ml) for 3 hr as indicated. The *x* axis represents the positions of the CpG sites relative to the transcription start site (+1); the *y* axis represents the percentage. Closed circle: methylated cytosine; Open circle: unmethylated cytosine.

### The TET1-mediated regulation of *Cxcl1* promoter methylation is important for the induction of *Cxcl1* gene

Next, we performed experiments to test if the TET1-mediated DNA methylation indeed contributes to the regulation of *Cxcl1* gene induction by luciferase assays. We first cloned a DNA fragment containing the Region#1 of the *Cxcl1* gene promoter into a luciferase reporter plasmid (pGL3-CXCL1). Transient transfection of this reporter plasmid in 293T cells showed a strong induction of luciferase activity in response to TNFα stimulation, indicating that the DNA fragment (−157 to +283) in this reporter plasmid indeed functions as the TNF-responsive promoter of *Cxcl1* gene (Fig. 5A).

**Figure 5.**
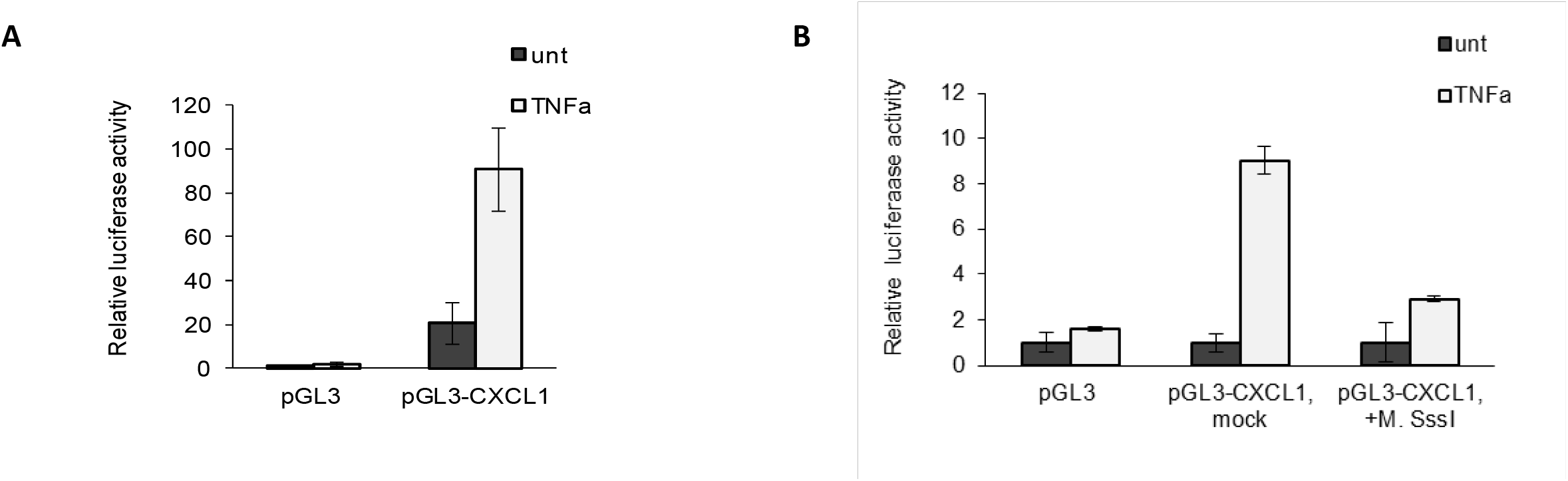
Analysis of the effect of DNA methylation of the *Cxcl1* gene promoter. (A) 293T cells were transfected with the empty vector control (pGL3) or the *Cxcl1* promoter containing plasmid (pCXCL1 promoter). Cells were then untreated with treated with human TNFα (10ng/ml) followed by luciferase assays. (B) Same as in (A) except 293T cells were transfected with the unmethylated *Cxcl1* promoter or the cytosine methylated *Cxcl1* promoter generated by *in vitro* treatments with or without the CpG methyltransferase M.SssI before plasmid ligation as indicated. Shown is the representative result of three independent experiments. Error bars represent SD.

To examine the role of CpG methylation in gene activation, the *Cxcl1* promoter fragment (nucleotide −157 to +283) was subjected to treatment with the CpG methyltransferase M.SssI *in vitro* as previously described ^56^. The M.SssI treated or mocked treated DNA fragments were then cloned into the pGL3 vector plasmid, followed by transient transfection into 293T cells together with the control Renilla plasmid. Consistent with the results from Fig. 5A, the mocked treated *Cxcl1* promoter plasmid showed strong luciferase reporter activity in response to TNFαstimulation. In contrast, the TNFα-induced reporter activity was significantly blocked in cells transfected with the M.SssI treated *Cxcl1* promoter plasmid (Fig. 5B). These studies suggest that the cytosine methylation in the Region #1-containing *Cxcl1* promoter results in the suppression of its promoter activity in response to TNFα treatment. Taken together, our results show that the TET1-mediated CpG demethylation of the Region #1 in the *Cxcl1* promoter may contribute at least in part to the induction of *Cxcl1* in response to LPS or TNFα stimulation.

### *Tet1* disruption resulted in defective neutrophil recruitment in the LPS-induced mouse model of acute lung injury

CXCL1 plays an important role in neutrophil recruitment. LPS can trigger acute lung injury with neutrophil emigration that contributes to the initiation of lung inflammation ^57–60^. We used the LPS-induced mouse model of acute lung injury to examine the physiological significance of the TET1 in neutrophil recruitment during lung inflammation. WT, *Tet1^+/−^* and *Tet1*^−/−^ mice were subjected to LPS challenge for 4 hr, and BAL fluid was collected and subjected to flow cytometry analysis. As shown in Fig. 6A, the recruitment of neutrophils was significantly reduced in the lungs of *Tet1^−/−^* mice. We also observed a modest increase of macrophages in the *Tet1^−/−^* mice, but the distribution of eosinophils was not affected.

**Figure 6.**
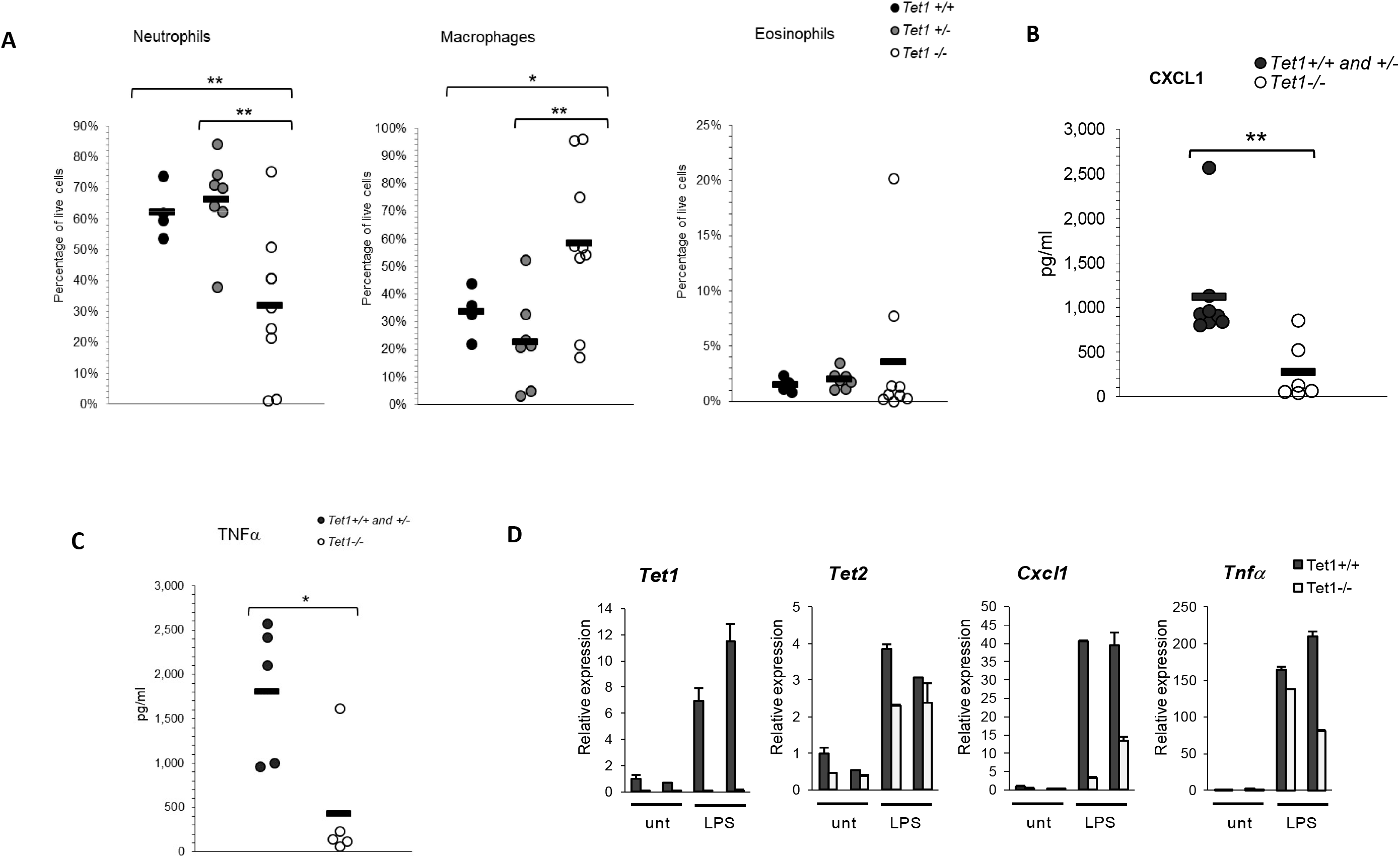
Defective neutrophil recuriment and *Cxcl1* production in BAL fluid of *Tet1*^−/−^ mice in LPS-induced acute lung injury. (A) WT and *Tet1*^−/−^ mice were intranasally injected with LPS (0.5mg/kg) for 4 hr, and Bronchoalveolar lavage (BAL) fluids were collected and analyzed by flow cytometry. Data were analyzed using the FlowJo software. Neutrophils were gated as CD45^+^CD11b^+^Ly6G^+^ cells, macrophages were identified as CD45^+^CD11c^+^SiglecF^+^ population of cells, and eosinophils were defined as CD45^+^CD11c^low/−^SiglecF^+^ cells. WT (n=4), *Tet1*^+/−^ (n=7), and *Tet1*^−/−^ (n=9). (B) The CXCL1 protein levels in BAL fluids from mice treated with LPS by intranasal injections for 4 hr were measured by ELISA assays. The *Tet1*^+/+^ and *Tet1*^+/−^ group (n=8); *Tet1*^−/−^ (n=6). (C) Same as in (B) except TNFα protein was measured by ELISA assays. The *Tet1*^+/+^ and *Tet1*^+/−^ group (n=5); *Tet1*^−/−^ (n=5). (D) Q-PCR analysis of RNA samples prepared from total cells obtained from the BAL fluid from two independent pairs of WT and *Tet1*^−/−^ mice treated intranasally with PBS or LPS (0.5 mg/kg) for 16 hr. *P* values were determined by non-paired *t*-test.

Next, we examined the overall CXCL1 protein levels in the BAL fluid by ELISA assays. Upon LPS stimulation, the total level of CXCL1 protein was significantly repressed in the BAL fluid obtained from *Tet1^−/−^* mice as compared to that from WT and *Tet1^+/−^* controls (Fig. 6B), which is consistently with the essential role of TET1 in the regulation of *Cxcl1* induction.

TNFα, a critical mediator of inflammation and tissue damage, is produced mainly by neutrophils and macrophages in response to LPS stimulation ^61^. To test if the defective neutrophil recruitment observed in *Tet1^−/−^* mice affects TNFα production in the BAL fluid, ELISA assays were performed to measure the levels of TNFα protein. As shown in Fig. 6C, the production of TNFα was significantly reduced in the BAL fluid of *Tet1*-deficient mice upon LPS challenge, consistent with the defective neutrophil recruitment in the absence of TET1.

We examined the mRNA levels of *Cxcl1* and *Tnfα* in the total cell population present in the BAL fluid following LPS challenge by Q-PCR analysis. Again, the induction of *Cxcl1* was defective in cells isolated from the *Tet1^−/−^* BAL fluids (Fig. 6D). *Tet1* disruption did not significantly affect the LPS-induced expression of *Tet2*. Consistent with the reduced percentage of neutrophils present in the total cell population of the BAL fluid of the LPS-treated *Tet1^−/−^* mice (Fig. 6A), the overall mRNA level of *Tnfα* was also repressed (Fig. 6D). These results support the *in vivo* functional role of TET1 in the specific regulation of the LPS-induced *Cxcl1* expression and neutrophil recruitment during acute lung injury.

## Discussion

The chemokine CXCL1 plays an important role in immune response and tumorgenesis. CXCL1 functions as a critical chemoattractant for neutrophils during lung inflammation. In cancer, it has been suggested that CXCL1 mediates tumor cell metastasis ^62^, and is important for the recruitment of myeloid-derived suppressor cells (MDSCs) into the tumor microenvironment ^63^. The promoter DNA methylation of *CXCL1* in human cancer cell lines has been observed ^24^, but its biological significance in tumorigenesis as well as the molecular basis involved is not clear. SETDB2, a histone lysine methyltransferase, was shown to be induced upon pathogenic infection and suppressed the induction of *Cxcl1* ^64^. The role of DNA methylation in the regulation of *Cxcl1* has not been understood. In this report, we provide both genetic and biochemical evidences to show that TET1 acts as a specific epigenetic regulator of *Cxcl1* by promoting the DNA demethylation and thus the transcriptional activation of *Cxcl1* in response to inflammatory stimuli such as LPS and TNFα.

A key functional property of TET proteins is their ability to promote DNA demethylation. The role of TET proteins in immune regulation is best exemplified by the finding that TET2 acts to restrict the production of inflammatory cytokines such as IL-6 and IL-1β in macrophages, and the loss of TET2 results in clonal hematopoiesis and accelerates atherosclerosis. In this case, it is proposed that TET2 functions as a transcriptional repressor to suppress inflammatory gene activation by recruiting HDACs, a mechanism that is independent of its activity in DNA demethylation ^38–40^. These findings have raised an interesting question on whether the TET-mediated DNA demethylation plays a role in immune regulation. Our studies described in this paper have demonstrated a clear role of TET1 in mediating the promoter DNA demethylation of *Cxcl1*. In contrast to TET2, which functions as a transcriptional repressor in immune gene activation, TET1 acts as an important activator in the induction of *Cxcl1* in response to inflammatory stimuli. Thus, TET1 and TET2 have opposing effects on immune gene activation.

There is also a clear difference between TET1 and TET2 in their specificity involved in gene regulation in macrophages. Gene profiling by microarray showed that the expression of about 450 genes was altered by more than a factor of 1.5 when compared WT and Tet2^−/−^ macrophage upon LPS stimulation ^40^. In contrast, the expression of only 48 genes was affected in *Tet1* deficient macrophages upon LPS stimulation, with the most significant effect on the induction of *Cxcl1*. Thus, TET1 appears to be more specific in immune gene regulation in macrophages. Interestingly, several other genes such as *Cspg4*, *Cpeb1*, *Htra4*, *Prg4*, and *B3gnt3* ^50–55^ that are known to play a role in cell migration and trafficking were found to be significantly altered by *Tet1* deletion. These results raise an interesting possibility that TET1 may have a broad role in the control of cell migration. The molecular basis for the observed specificity of TET1 in macrophage gene regulation remains to be unclear. One possibility is the presence of a very low amount of TET1 protein in macrophages. In fact, although we could clearly detect *Tet1* mRNA by Q-PCR analysis (Fig. 1A), we were unable to detect TET1 protein in BMDMs or MEFs with different sources of anti-TET1 antibodies (unpublished observation). Thus, it is likely that the cell-type specificity of an epigenetic regulator such as TET1 can be achieved by modulating its level of protein expression.

Consistent with the well-documented importance of CXCL1 in neutrophil recruitment, we showed that the presence of neutrophils in the BAL fluid was significantly suppressed in *Tet1*^−/−^ mice upon LPS challenge as compared to the WT or *Tet1^+/−^* control mice. In contrast, the distribution of eosinophils in the BAL fluid was not altered by *Tet1* disruption. Interestingly, we observed a modest increase of macrophages in the *Tet1*^−/−^ BAL fluid. Despite the increased resident macrophages which are known to be the major CXCL1-producing cells in the *Tet1*^−/−^ BAL fluid, the level of CXCL1remained to be suppressed, a phenomenon consistent with the conclusion that macrophages with *Tet1* disruption are defective in producing CXCL1. In conclusion, our studies have identified an epigenetic mechanism that regulates the CXCL1-mediated neutrophil recruitment during lung inflammation.

## Acknowledgements

We thank the UCLA flow cytometry core facility and the UCLA Technology Center for Genomics and Bioinformatics (TCGB) core facility for technical support. Supported by funds from the UCLA Division of Hematology-Oncology (K.S.), by grants from the Foundation of Sichuan Provincial Science and Technology Program (#2019YFH0158 and #2019YFH0147 to J.Z.), and by Project for Disciplines of Excellence Award from West China Hospital, Sichuan University (ZYJC18016 to J.Z. and J.W.).

## Authorship contributions

J.Z., J.W. and K.S. directed the project and designed experiments. K.S., K.M.Y., J.Z., J.W., and R.W.S. wrote the manuscript. K.M.Y. executed most of the experiments. R.W.S. analyzed and interpreted RNA-seq data and performed statistical analysis. B.L., C.A.H., C.N., H.R.L., M.S.L., and J.W. participated in Q-PCR analysis and animal studies.

## Supplementary Data

### Material and methods

#### Cell culture

Mouse bone marrow-derived dendritic cells (BMDC): Bone marrow was collected and red blood cells lysed similarly as for BMDM indicated above. Then 2 ×10^6^ cells were seeded per 35 mm well with 1.5 ml of complete RPMI containing 10% FBS, 1% penicillin-streptomycin and 20 ng/ml recombinant murine GM-CSF (PeproTech). On day 2 and day 4, most of the media was carefully removed to avoid disturbing the non-adherent cells, and replaced with 1.5 ml of fresh, complete RPMI. On day 6, the non-adherent cells were collected and reseeded onto fresh plates. On day 7, cells were used for treatment assays.

Peritoneal macrophages: Peritoneal cell exudate (PEC) was harvested by careful injection of 5 ml of cold serum-free RPMI into the peritoneal cavity of ethanized mice via a 25 G needle and 5 ml syringe. Care was placed to avoid puncturing major organs and blood vessels. The peritoneum was then gently massaged for a few seconds to loosen any attached cells into the RPMI solution. Then using the same needle and syringe, the injected RPMI was carefully collected. The procedure was repeated with another 5 ml of RPMI, for a total volume of 10 ml.

Samples with blood were discarded. PEC was washed once with PBS, then resuspended in complete RPMI containing 10% FBS and 1% penicillin-streptomycin. Cells were seeded onto 12-well plates and placed in a 37°C tissue-culture incubator. The next day, non-adherent cells were washed off with PBS. Fresh complete media was added to the adherent cells (i.e. peritoneal macrophages). After resting cells for at least an hour in the incubator, cells were then used for experiments.

---

**Figure S1.**
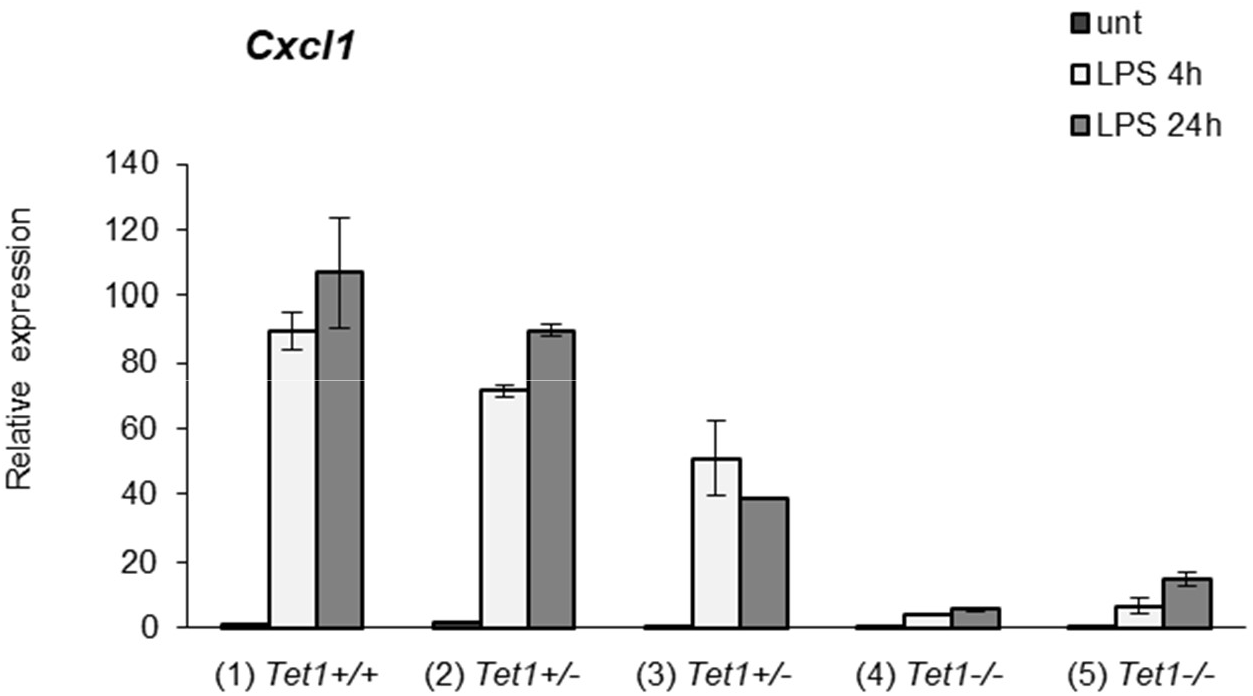
Gene-dosage effect of *Tet1* on Cxcl1 induction in response to LPS in BMDMs. Q-PCR analysis of *Cxcl1* induction in BMDMs prepared from WT [(1)], *Tet1*^+/−^ [(2), (3)], *Tet1*^−/−^ [(4), (5)] treated with LPS (10ng/ml) for 4 hr and 24 hr as indicated. Error bars represent SD.

**Figure S2.**
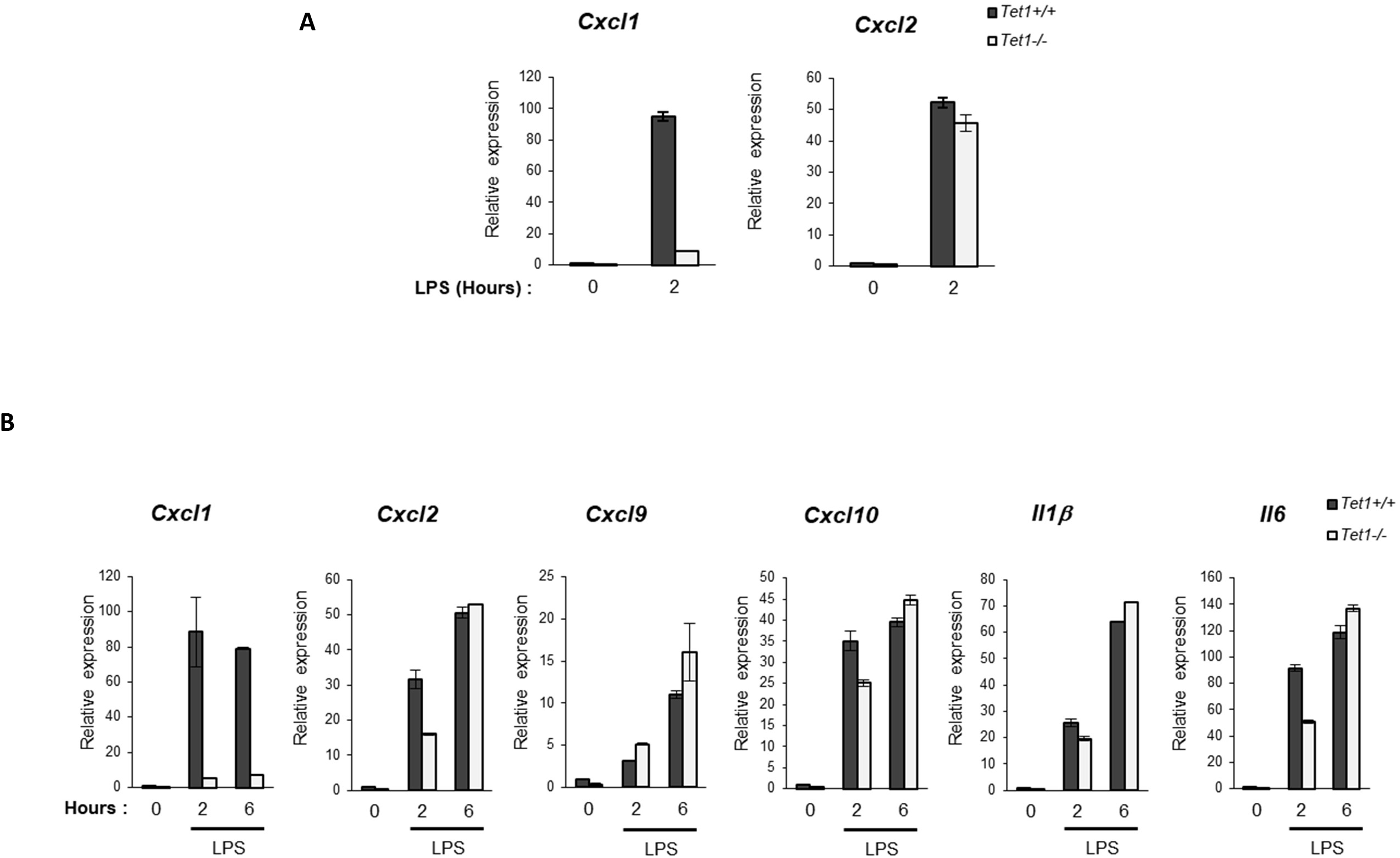
The effect of TET1 on Cxcl1 induction in peritoneal macrophages and BMDCs. (A) Q-PCR analysis of *Cxcl1* and *Cxcl2* induction in primary peritoneal macrophages of WT and *Tet1*^−/−^ mice untreated or treated with LPS (10ng/ml) for 2 hr. (B) Same as in (A) except RNA samples from BMDCs were used. Error bars represent SD.

**Figure S3.**
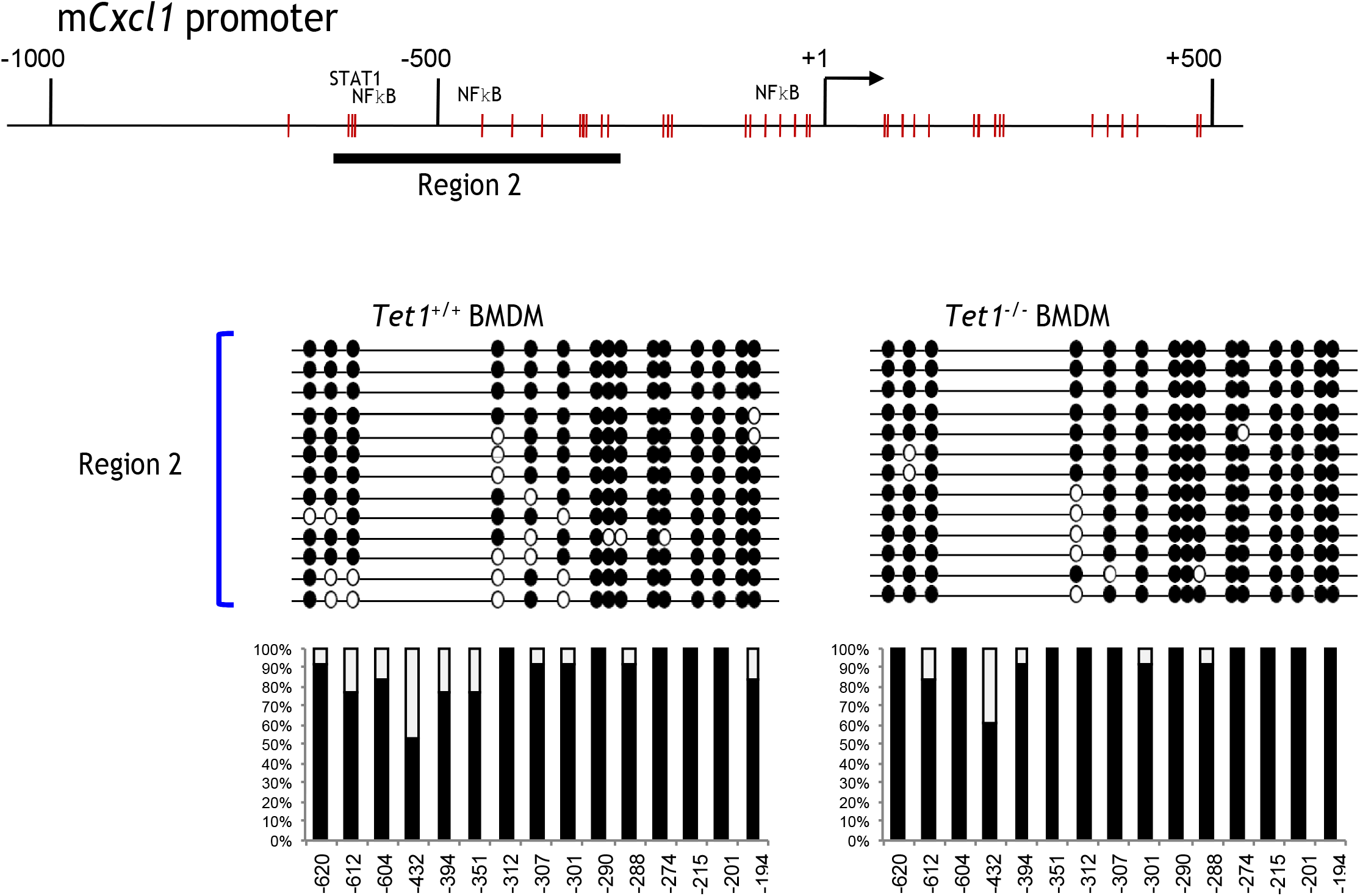
Analysis of DNA methylation in the region #2 of the *Cxcl1* promoter in BMDMs. Methylation analysis of the *Cxcl1* promoter region#2 was performed by bisulfite conversion of genomic DNA from WT and *Tet1*^−/−^ BMDMs. The *x* axis represents the positions of the CpG sites relative to the transcription start site (□1); the *y* axis represents the percentage. Closed circle: methylated cytosine; Open circle: unmethylated cytosine.

**Figure S4.**
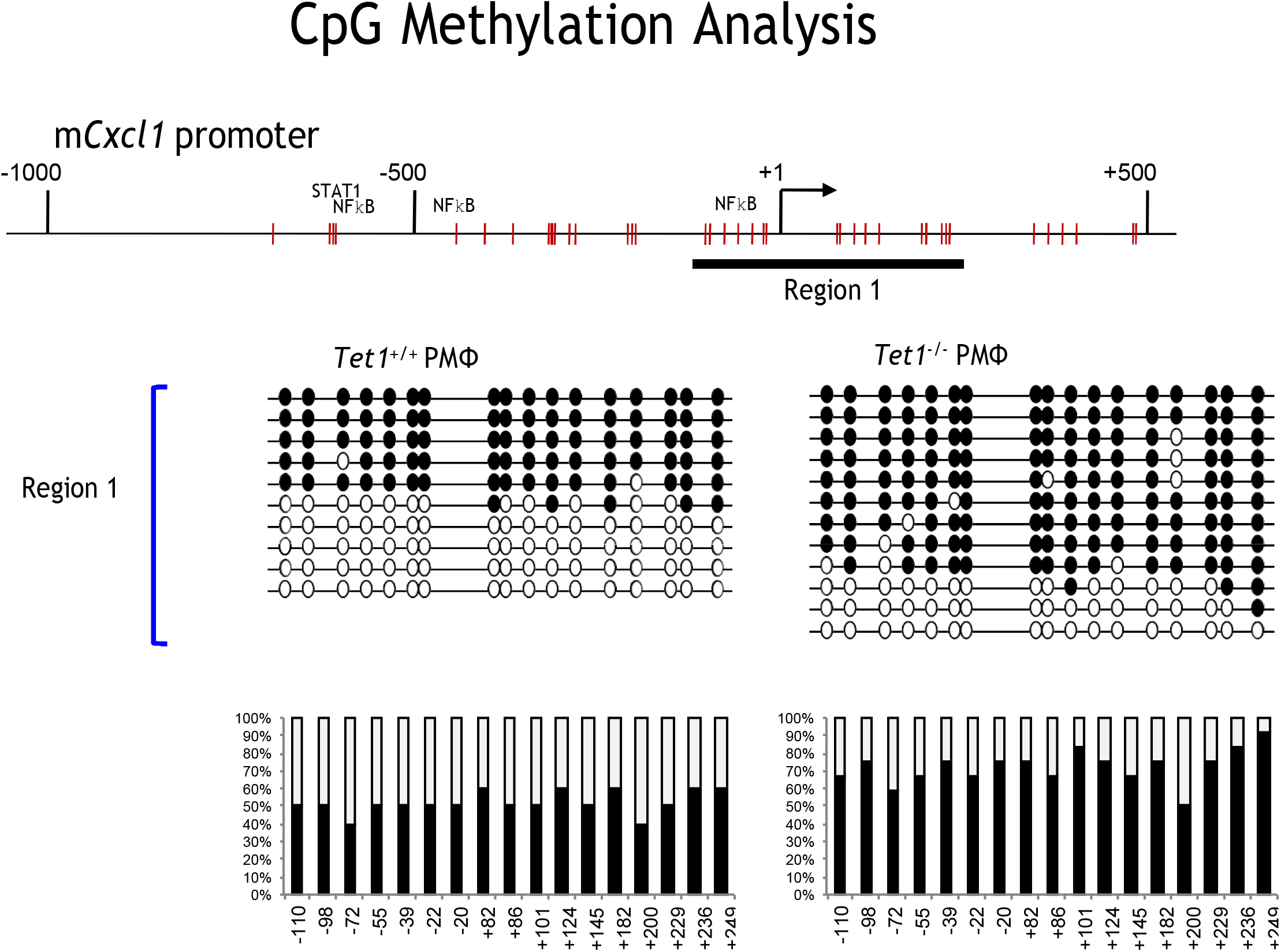
TET1 regulates the DNA methylation of the *Cxcl1* promoter in peritoneal macrophages. Methylation analysis of the *Cxcl1* promoter region#1 was performed by bisulfite conversion of genomic DNA from WT and *Tet1*^−/−^ peritoneal macrophages. The *x* axis represents the positions of the CpG sites relative to the transcription start site (□1); the *y* axis represents the percentage. Closed circle: methylated cytosine; Open circle: unmethylated cytosine.

**Figure S5.**
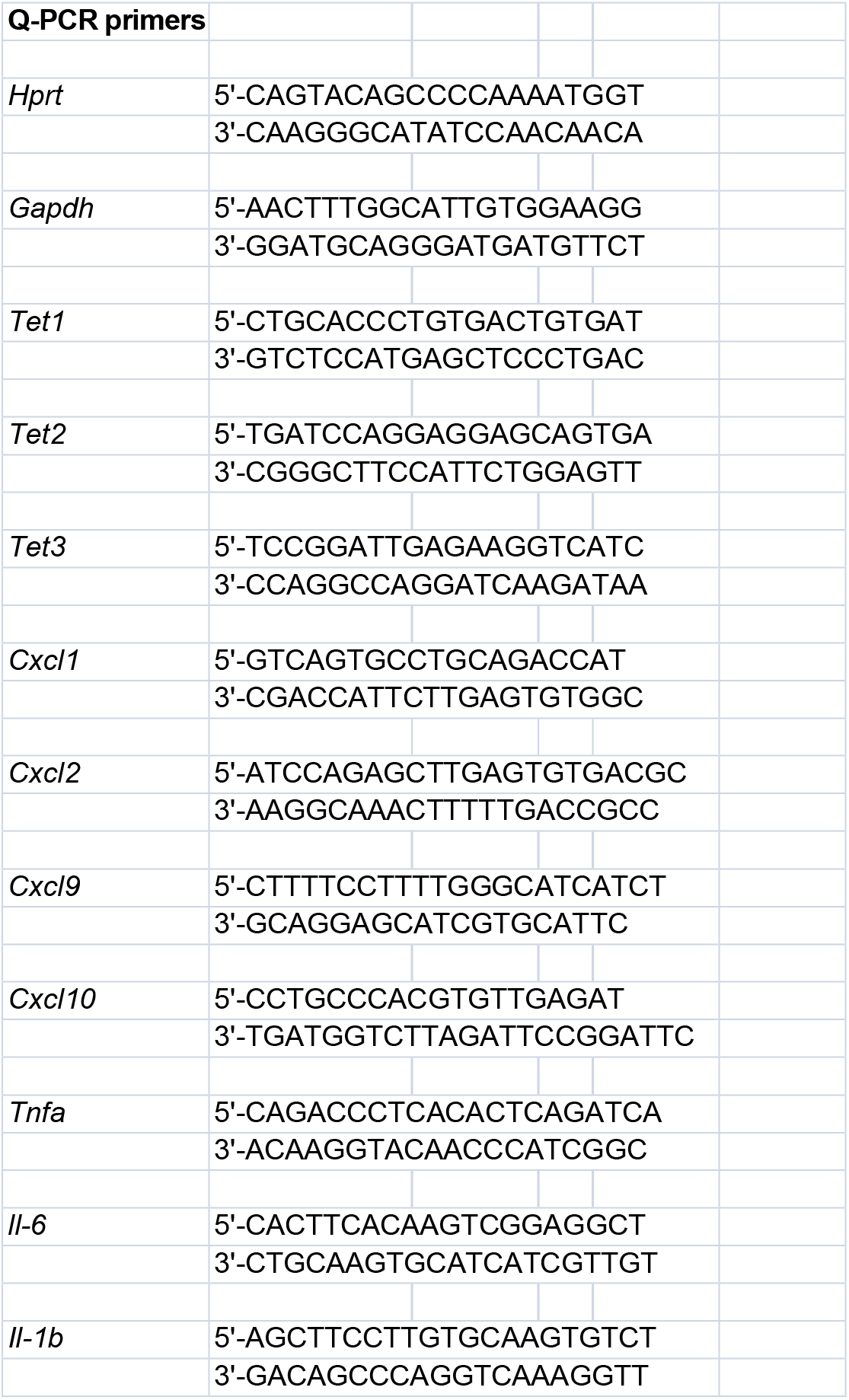
List of Q-PCR primers. See detailed description in the figure.

**Figure S6.**
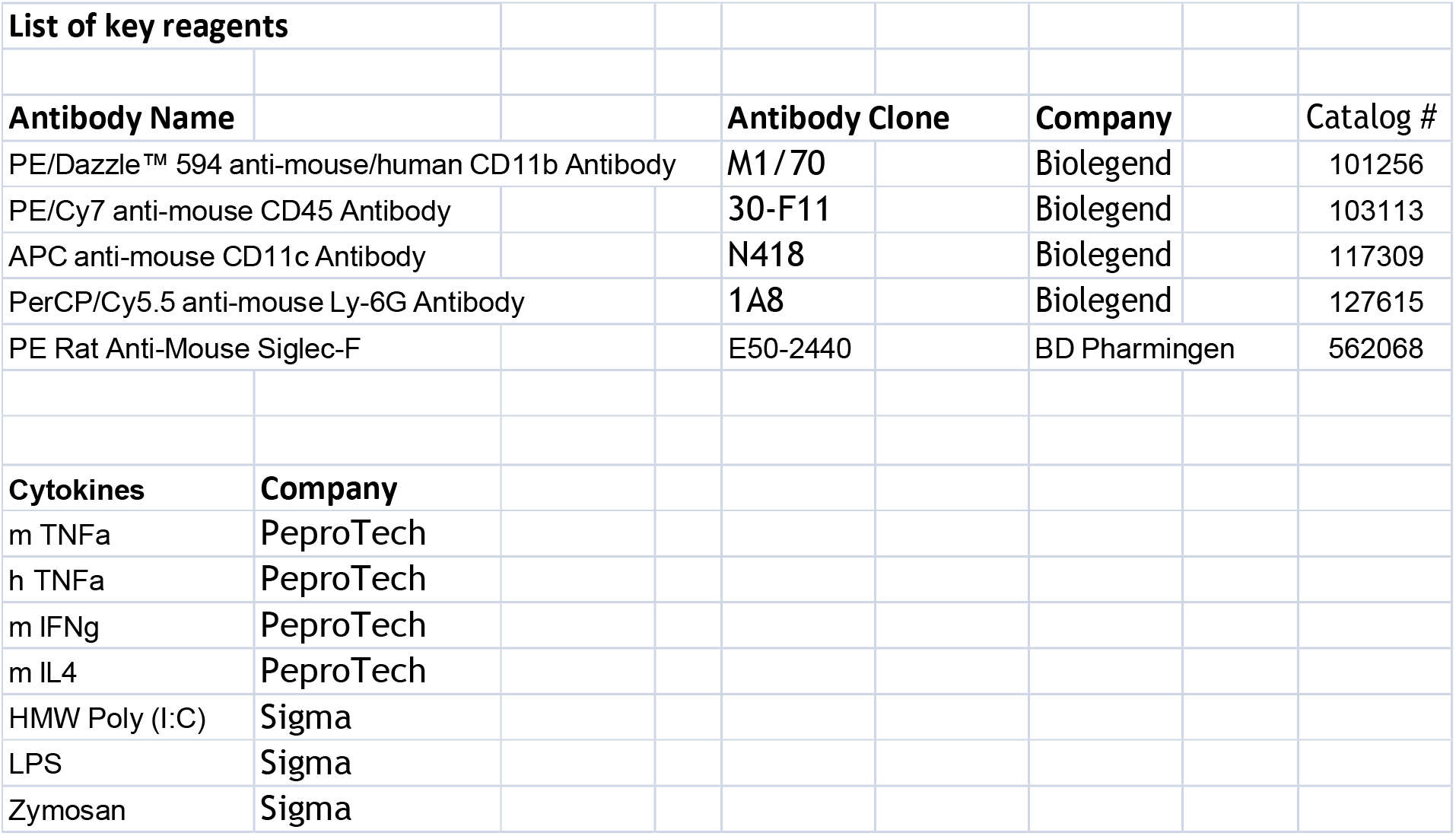
List of key reagents. See detailed description in the figure.

